# IBD-based estimation of X chromosome effective population size with application to sex-specific demographic history

**DOI:** 10.1101/2022.07.06.499007

**Authors:** Ruoyi Cai, Brian L. Browning, Sharon R. Browning

## Abstract

The effective size of a population (*N*_*e*_) in the recent past can be estimated through analysis of identity-by-descent (IBD) segments. Several methods have been developed for estimating *N*_*e*_ from autosomal IBD segments, but no such effort has been made with X chromosome IBD segments. In this work, we propose a method to estimate the X chromosome effective population size from X chromosome IBD segments. We show how to use the estimated autosome *N*_*e*_ and X chromosome *N*_*e*_ to estimate female and male effective population sizes. We demonstrate the accuracy of our autosome and X chromosome *N*_*e*_ estimation with simulated data. We find that estimated female and male effective population sizes generally reflect the simulated sex-specific effective population sizes across the past 100 generations, but that short-term differences between the estimated sex-specific *N*_*e*_ across tens of generations may not reliably indicate true sex-specific differences. We analyzed the effective size of populations represented by samples of sequenced UK White British and UK Indian individuals from the UK Biobank.

## Introduction

The effective size of a population (*N*_*e*_) is defined as the number of breeding individuals in an idealized randomly mating population that has the same expected value of a parameter of interest as the actual population under consideration (Wright 1931). The effective population size is a fundamental parameter in population genetics because it determines the strength of genetic drift and the efficacy of evolutionary forces such as mutation, selection, and migration (Charlesworth 2009). Previous studies have demonstrated that estimates of recent effective population size can reveal aspects of a population’s demographic history, such as past population growth or bottleneck events (Browning and Browning 2015; Browning *et al*. 2018).

Identity-by-descent (IBD) segments can be used to estimate effective population size in the recent past. IBD segments are haplotypes which two or more individuals have inherited from a common ancestor. IBD segments end at positions where crossovers have occurred in the meioses between the common ancestor and the descendant individuals. IBD segments for which the common ancestor is in the distant past tend to be shorter than IBD segments from a recent common ancestor because there are more meioses since the common ancestor on which crossovers can occur. The autosomes and the X chromosome are both subject to recombination, making them both amenable to IBD segment analysis (Buffalo *et al*. 2016; Henden *et al*. 2016). Previous studies have developed methods for estimating recent effective population size from autosomal IBD segments (Palamara *et al*. 2012; Browning and Browning 2015; Browning *et al*. 2018). However, no such effort has been made with X chromosome IBD segments.

The autosomes are equally influenced by female and male demographic processes. In contrast, the X chromosome is influenced more strongly by female demographic processes than by male demographic processes. This is because females have two copies of the X chromosome, while males have only one. Thus, comparison of statistics from the X chromosome and from the autosome can be used to estimate sex-specific parameters such as female and male effective population sizes (Hammer *et al*. 2008; SÉgurel *et al*. 2008; Bustamante and Ramachandran 2009; Keinan *et al*. 2009; Bryc *et al*. 2010; Heyer *et al*. 2012; Goldberg and Rosenberg 2015; Buffalo *et al*. 2016; Shringarpure *et al*. 2016; Clemente *et al*. 2018; Musharoff *et al*. 2019).

The standard Wright-Fisher model used to define the effective population size assumes equal numbers of breeding females and males. To define female and male effective population sizes, we consider an idealized Wright-Fisher population modified to allow for different numbers of females and males. The female and male effective sizes (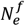 and 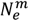) of a study population are the number of females and males in an idealized Wright-Fisher population that would give the same rates of IBD on autosomes and sex chromosomes as the study population (Wang *et al*. 2016).

In this work, we develop an IBD-based method to estimate the X chromosome effective population size in each generation in recent history. We show that estimated X chromosome *N*_*e*_ can be combined with estimated autosome *N*_*e*_ to estimate the female and male effective population sizes over time. This application of the X chromosome *N*_*e*_ provides a useful complement to previous methods that give only a single time-averaged estimate for the sex-specific effective sizes in human populations (Hammer *et al*. 2008; Emery *et al*. 2010; Clemente *et al*. 2018). Through simulation studies, we validate the theoretical relationship between the autosome and X chromosome *N*_*e*_, and we show that our method can accurately estimate the autosome and X chromosome *N*_*e*_ in simulated populations. We investigate the application of X chromosome *N*_*e*_ to estimate sex-specific *N*_*e*_ and show that pronounced differences in the estimated female and male effective population sizes over many generations are reasonably accurate, but confidence intervals do not always cover the true values, and short-term observed differences in the estimated effective population sizes may not represent true differences. We use our method to infer autosome and X chromosome *N*_*e*_, as well as sex-specific *N*_*e*_, for several human populations.

## Methods

### Probability modelling for the X chromosome

All meioses from mothers transmit an X chromosome, while half of meioses from fathers transmit an X chromosome. Over many generations, approximately two-thirds of meioses from parents to offspring that transmit an X chromosome will be from females, as we show below. However, the actual proportion of meioses transmitting an X chromosome that are from females in recent generations depends on the proportion of females in these generations. For example, if the sample consists entirely of females (who receive an X chromosome from each parent), half of the meioses in the most recent generation that transmit an X chromosome are from females. If the sample is made up solely of males (who receive only one X chromosome, inherited from the mother), all the meioses that directly contribute the X chromosomes in the sample are from females. If half the sampled individuals are female and half are male, then 2/3 of the meioses in the previous generation that contribute the X chromosomes in the sample will be from females. We show in the next paragraph that when half of the samples are female the expected value for this ratio will be 2/3 in all prior generations, regardless of sex-specific demographic forces.

Consider the lineage of a randomly selected haplotype at a point in the genome. Let *p*_*g*_ be the probability that the ancestral haplotype at generation *g* before present is carried by a female, where *g* = 0 corresponds to the generation of the sampled individuals, *g* = 1 to the generation of their parents, and so on. A given haplotype carried by a female has a 50% probability that its parent haplotype is carried by a female, while a given haplotype carried by a male always has its parent haplotype carried by a female. Thus, for *g* ≥ 0,

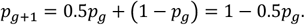

If *p*_*g*_ = 2/3, then *p*_*g*+1_ = 2/3. If the sampled haplotype is randomly chosen from a set of individuals with equal numbers of females and males, then the sampled haplotype has probability *p*_0_ = 2/3 of being carried by a female, and as a result *p*_*g*_ = 2/3 for all *g*. More generally, it can be shown by mathematical induction that

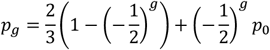

which converges to 2/3 for large *g*. This equation is similar to equations for X chromosome admixture proportions and for X chromosome allele frequencies in a particular breeding system (Goldberg and Rosenberg 2015; Rosenberg 2016). In what follows, we assume that *p*_*g*_ = 2/3 for all *g*.

Consider IBD sharing on the X chromosome resulting from a common ancestor living *g* generations ago. Let *F* be the number of female meioses out of the 2*g* meioses in the path of inheritance. Then *F* follows a binomial(2*g*, 2/3) distribution. Assuming Haldane’s model (Haldane 1919), the length of the IBD segment is exponentially distributed with rate 3*F*/2 per Morgan, as crossovers that end an IBD segment happen only in female meiosis on the X chromosome (which comprise 2/3 of the meioses) and the female recombination rate is 3/2 per Morgan on the X chromosome when using sex-averaged genetic distances. As an approximation, we model the probability distribution of the length of an X chromosome IBD segment resulting from a coalescence event occurring *g* generations ago as an exponential random variable with rate 2*g* per Morgan, which is the same model that we use for estimating IBD-based effective population size in autosomal data (Browning and Browning 2015). We compared the simulated distribution of draws from an exponential(3*F*/2) distribution with *F* drawn from a binomial(2*g*, 2/3) distribution to the exponential(2*g*) distribution for different values of *g* (Figure S1). The result shows that the exponential(2*g*) model approximates the distribution of IBD length on the X chromosome very well for *g* > 1.

### IBD-based estimation of X chromosome effective population size

Our method for IBD-based estimation of X chromosome effective population size history is based on IBDNe which was designed to estimate recent effective population size from autosomal IBD segments (Browning and Browning 2015). The IBDNe method calculates the expected length distribution of IBD segments exceeding a given length threshold (2 cM by default) for a given effective population size history. It finds the effective population size history that equates the observed and expected IBD length distributions using an iterative scheme. IBDNe applies smoothing over intervals of eight generations to avoid overfitting. Although IBDNe was designed for autosomal data, we show that it can also be used with X chromosome data, with some adjustments to the analysis procedure that we describe in the following paragraphs.

The first adjustment ensures proper inference of IBD segments on the X chromosome by encoding male X chromosome genotypes as haploid. Male X chromosome genotypes are frequently coded as homozygous diploid genotypes rather than haploid genotypes, which typically results in duplicate reported IBD segments when using IBD detection methods designed for autosomal data. The hap-ibd program (Zhou *et al*. 2020) correctly analyzes chromosome X data with haploid male genotypes. Alternatively, if males are coded as homozygous diploid on the X chromosome, we can exclude IBD pairs involving the second haplotype of males after running hap-ibd on the diploid-coded data. We exclude the pseudo-autosomal regions and regions outside the genetic map from analysis.

The second adjustment is to use a sex-averaged genetic map, as we also do for the autosomes. X chromosomes transmitted from females are subject to recombination, while X chromosomes transmitted from males are not (except for the pseudo-autosomal regions which we exclude from all analyses). Genetic maps, such as the HapMap map (International Hapmap Consortium 2007) and the deCODE map (Halldorsson *et al*. 2019), typically report the female-specific recombination map for the X chromosome. Since an average of 2/3 of meioses transmitting an X chromosome are from females, the sex-averaged X chromosome recombination map can be obtained by multiplying female-specific genetic distances by 2/3. For example, a region with length 3 cM on the female-specific map has length 2 cM on the sex-averaged map. Equivalently, the sex-averaged recombination rates can be obtained by multiplying female-specific recombination rates by 2/3. For example, an X chromosome region with female-specific recombination rate of 3 × 10^−8^ per base pair per generation has a sex-averaged recombination rate of 2 × 10^−8^ per base pair per generation.

The third adjustment ensures equal numbers of sampled females and males. If the sample is unbalanced, we remove some randomly selected females or males to obtain equal numbers of females and males. Consequently, *p*_0_, the proportion of sampled X chromosome haplotypes carried by females, is 2/3, and hence *p*_*g*_, the probability that the ancestral haplotype of a sampled X chromosome haplotype *g* generations before the present is carried by a female is always 2/3 (see the preceding “Probability Modelling for the X chromosome” section).

The fourth adjustment modifies the IBDNe “npairs” parameter to be equal to the number of analyzed haplotype pairs. By default, IBDNe assumes that each individual contributes two haplotypes to the analysis, and that all cross-individual pairs are analyzed, resulting in (2*n*)(2*n* − 2)/2 haplotype pairs when there are *n* individuals. On the X chromosome, with *n*_*f*_ females and *n*_*m*_ males, the number of haplotype pairs is

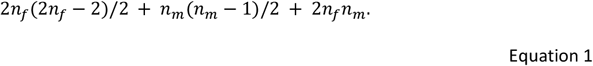

We set the IBDNe “npairs” parameter to the value in equation 1 when analyzing X chromosome data, after adjusting it for removal of close relative pairs as described below.

The fifth adjustment is to manually remove detected IBD segments corresponding to close relatives (parent-offspring and siblings). By default, IBDNe identifies the close relatives using the input IBD segments, removes the IBD segments between them, and adjusts the “npairs” parameter to account for the removed sample pairs. However, this strategy does not work for the X chromosome because one cannot reliably detect close relatives using only X chromosome data. We thus turn off IBDNe’s filtering of close relatives by setting “filtersamples=false”. We can identify close relatives based on autosomal data or from a pedigree file if available, and then remove IBD segments for these pairs and update the “npairs” parameter accordingly in the chromosome X analysis. Removing only IBD segments between the related pairs rather than completely removing one individual from each pair of relatives reduces the loss of information from the data.

The sixth adjustment enables calculation of confidence intervals. IBDNe obtains confidence intervals for the estimated effective population sizes by bootstrapping over chromosomes. We thus divide the X chromosome into six pieces of equal cM length and treat these as separate “chromosomes” in the analysis with IBDNe.

### From X chromosome effective population size to sex-specific effective population size

We next describe how the estimated X chromosome effective population size can be used in conjunction with the estimated autosome effective population size to estimate female and male effective population sizes. We will write 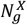 and 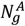 for the X chromosome and autosomal effective population sizes at generation *g*. And we will write 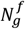 and 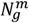 for the female and male effective population sizes at generation *g*, which can be derived from the X chromosome and autosomal effective population sizes as described below.

The IBD-based effective population size (for autosomes or X chromosome) is defined in terms of the conditional coalescence probability for a Wright-Fisher population. We first consider autosomes. For a randomly selected pair of haplotypes, let *G* be the number of generations before present that the haplotypes coalesce. Conditional on the haplotypes not coalescing by generation *g* − 1 before present, the ancestral haplotypes are distinct at that generation. For them to coalesce at generation *g* before present, their two parental haplotypes at generation *g* must be the same haplotype. If the diploid autosomal effective population size is 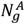 at *g* generations before present, there are 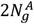 autosomal haplotypes available, and the probability that the two parental autosomal haplotypes are the same is thus 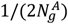. That is, 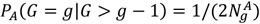. Thus, if we know the value of *P*_*A*_(*G* = *g*|*G* > *g* − 1), then we can obtain the effective population size *g* generations before present:

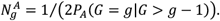

On the X chromosome, the conditional coalescence probability can be obtained by considering that 2/3 of meioses are from female parents, while 1/3 are from male parents. For coalescence to occur, both haplotypes’ parent haplotypes must be the same. This means both haplotypes must be inherited from parents that have the same sex, and the second haplotype must have the same parental haplotype as that of the first (within that sex). Thus,

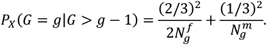

And hence,

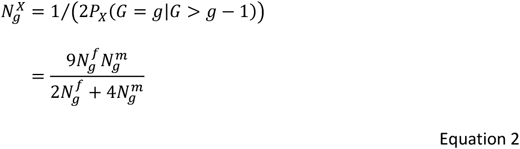

For comparison, on the autosomes, by the same reasoning but with half of the meioses from each sex and with diploid males,

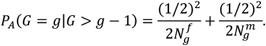

And hence,

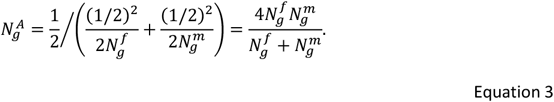

Equation 3 From Equations 2 and 3, the ratio of X to autosomal effective population size, which we denote as *α*, is

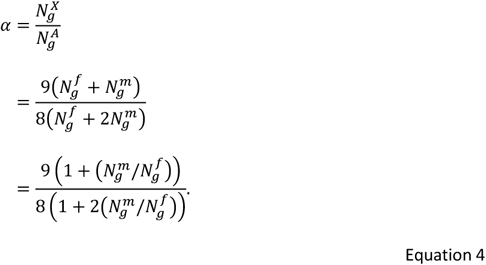

Thus 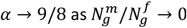, and 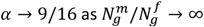. Since *α* is a decreasing function of 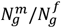, the ratio of X to autosomal effective population size satisfies 9/16 < *α* < 9/8.

With algebra, it can be shown that Equations 2 and 3 imply that

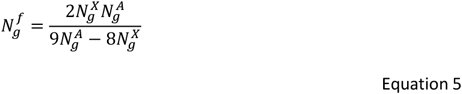

and

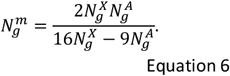

Thus, given estimates of 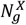 and 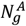, one can use Equations 5 and 6 to obtain estimates for 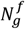 and 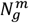. These are standard equations for estimating sex-specific effective population sizes based on 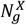 and 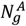 (Wright 1969; Hartl and Clark 1997), although usually presented in the context of constant effective population sizes across time.

According to Equation 4, the allowable range of X chromosome *N*_*e*_ is between 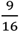 and 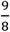 of the autosome *N*_*e*_ at each generation. When the X chromosome *N*_*e*_ is overestimated or the autosome *N*_*e*_ is underestimated, the estimated female effective population size (Equation 5) can be negative. Similarly, underestimation of the X chromosome *N*_*e*_ or overestimation of the autosome *N*_*e*_ can result in a negative estimate of the male effective population size (Equation 6).

### Analysis pipeline

We start with phased sequence data (using true phase for the simulated data, and inferred phase for real data), with males coded as haploid on the X chromosome. We use hap-ibd (Zhou *et al*. 2020) to infer IBD segments. For hap-ibd analysis on sequence data, we set the minimum seed length to 0.5 cM and the minimum extension length to 0.2 cM. The relatively small minimum seed length and minimum extension length increase power to detect short IBD segments. We exclude rare variants by setting the minimum minor allele count filter to 100 because these lower frequency variants are less informative, have less accurate phasing, and may be recent mutations.

We then run IBDNe on the detected IBD segments, with one analysis for autosomes and a separate analysis for the X chromosome. The genetic map file for the analysis is assumed to be a sex-averaged map. For the X chromosome, this means multiplying cM positions in the female-specific map by 2/3. When applying IBDNe on the simulated data (autosomes or X), we set “filtersamples=false” and “gmin=1” because the chromosomes are simulated independently so that a pair of individuals can share ancestry one generation back (i.e. be siblings) on one chromosome without such sharing occurring on other chromosomes. These settings tell IBDNe not to look for and remove close relatives, and to model IBD from shared ancestry starting from one generation before present.

Real data often has an excess of close relatives due to the sampling scheme. Thus, in the analysis of real autosomal data we allow IBDNe to detect and remove close relatives, which is the default behavior. The X chromosome on its own is not sufficient to detect close relatives, so we use either available pedigree information or the close relative pairs identified by IBDNe in the autosomal analysis to manually remove IBD segments from close relatives in the X chromosome data. We then update the “npairs” parameter accordingly (see below), and set “filtersamples=false” in the X chromosome IBDNe analysis.

In the IBDNe analysis of the X chromosome data, we set the “npairs” parameter equal to the number of haplotype pairs for which IBD segments could be present (i.e., all pairs except for those for which we have explicitly removed IBD segments). We first calculate the number of haplotype pairs based on the numbers of females and males in the sample, using Equation 1. We then adjust the number of haplotype pairs to account for the number of close relative pairs that were removed. Removing a male-male pair decreases the count by 1. Removing a male-female pair decreases the count by 2. Removing a female-female pair decreases the count by 4.

In order for IBDNe to obtain bootstrap confidence intervals for the X chromosome effective population size estimates, we divide the X chromosome into six pieces of equal cM length. We recode the chromosome field of the IBD segment file using integer values between 1 and 6 according to the location of the IBD segments. IBD segments that cross more than one of these “chromosomes” are split into subsegments at the boundaries of the corresponding “chromosomes”.

After obtaining the X chromosome and autosomal effective population sizes, we estimate the female and male effective population sizes using Equations 5 and 6. We obtain bootstrap values for these estimates by taking pairs of bootstrap values from the X and autosomal analyses. For example, for the *n*-th bootstrap value of the female effective population size at generation *g*, we take the *n*-th bootstrap value for the X chromosome effective population size at generation *g* and the *n*-th bootstrap value for the autosomal effective population size at generation *g*, and apply Equation 5. After obtaining all the bootstrap values, we use the 2.5^th^ and 97.5^th^ percentile to obtain an approximate 95% confidence interval for the female (for example) effective population size at generation *g*.

### Simulation study

We conducted a simulation study to evaluate the performance of our method. We used SLiM, a forward simulator (Haller and Messer 2019), to simulate the demographic history for the most recent 5000 generations, and we used msprime, a coalescent simulator, to complete the simulation back to full coalescence of the sample (Kelleher *et al*. 2016; Haller *et al*. 2019). For all scenarios, we simulated data for 30 autosomes of length 100 Mb and an X chromosome of length 180 Mb.

For the simulation in SLiM, we used a mutation rate of 10^−8^ per base pair per generation, and a recombination rate of 10^−8^ per base pair per meiosis on the autosomes and for female meioses on the X chromosome. We set the gene conversion initiation rate to 2 × 10^−8^ per base pair per meiosis on the autosomes and per female meiosis on the X chromosome, with mean gene conversion tract length of 300 bp. We simulated populations with equal sex ratio, and we also simulated populations with 20% females, 40% females, 60% females, and 80% females. We used the same total effective population size 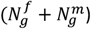 in each simulation. We sampled 50,000 individuals comprising 25,000 females and 25,000 males for each analysis.

We simulated data under a demographic model with a four-stage exponentially growing population with increased growth rate over time, which we call the “UK-like” scenario because it approximates the demographic history of the UK population (Browning and Browning 2013). The forward simulation of this model in SLiM starts from 5000 generations ago with an initial size of 3000. At 300 generations ago, this population starts to grow at an exponential rate of 1.4% per generation. At 60 generations ago, the rate of the exponential growth increases to 6%. In the most recent 10 generations, the growth rate further increases to 25%, and the population size reaches around 21 million at the time of sampling.

The part of the simulation conducted in msprime corresponds to the demographic history earlier than 5,000 generations ago that has a constant population size of 3,000. When completing the simulations with msprime, we did not apply gene conversion or sex-specific population size. In the msprime simulations, mutation occurred at a rate of 10^−8^ per base pair per generation, and recombination at a rate of 10^−8^ per base pair per meiosis. There is no differentiation between female and male meioses in the generations prior to the starting generation of the forward simulation because msprime does not have sex-specific functionality. This lack of sex-specific treatment in the period more than 5000 generations ago will affect the level of variation in the data but will not affect the distribution of IBD segments of length > 2 cM (the segments analyzed by IBDNe) because such segments typically have ancestry within the past three hundred generations.

### Analysis of human populations

We applied the above analysis pipeline on whole genome sequence data from the UK Biobank (Bycroft *et al*. 2018). The UK Biobank is a large-scale biomedical database that contains in-depth genetic, physical and health data collected between 2006 and 2010 on half a million UK participants aged between 40 and 69 (Fry *et al*. 2017). The UK Biobank whole-genome sequencing (WGS) consortium recently released high-coverage whole genome sequence data for 200,031 study participants (Halldorsson *et al*. 2022). We used Beagle 5.4 (Browning *et al*. 2021) and our previously-described UK Biobank phasing pipeline (Browning and Browning 2023) to phase the 200,031 genomes on the UK Biobank Research Analysis Platform. We used 2 cM as the IBD length threshold and estimated effective population size over the past 100 generations. We used the deCODE map in the analyses (Halldorsson *et al*. 2019).

We first analyzed the White British participants, who form the largest ethnic group in the UK Biobank. The UK Biobank sequence data include 91,532 White British females and 75,298 White British males. Since IBDNe has limits on the number of IBD segments that it can process, we randomly selected 5,000 White British females and 5,000 White British males for estimation of the autosome *N*_*e*_. For analysis of the X chromosome *N*_*e*_, we removed 16,234 randomly selected females to ensure equal numbers of females and males in the sample. For the autosomes, IBDNe removed close relatives using default settings. For the X chromosome, we used the UK Biobank’s kinship estimates to identify and remove IBD segments from sibling pairs and parent-offspring pairs (Bycroft *et al*. 2018).

The UK Biobank also includes participants from several ethnic minority groups including Black British, Indian, Pakistani, Asian, and Bangladeshi. Among these, we chose to analyze the effective population size of the Indian group, which has a large sample size, although we note that this group contains considerable diversity. There are sequence data for 1258 males and 1293 females with Indian ancestry in the UK Biobank. We removed 35 randomly selected females to achieve equal numbers of females and males for the subsequent analysis. By default, IBDNe automatically removes IBD segments from pairs of related individuals and generates a list of these related pairs. We manually removed X chromosome IBD segments for these pairs of related individuals prior to estimation of X chromosome *N*_*e*_ using IBDNe.

## Results

### Simulation study

We checked that the distribution of X chromosome IBD segments in female-female haplotype pairs, that in female-male haplotype pairs, and that in male-male haplotype pairs are consistent in all simulations (Figure S2). We compared the estimated X chromosome *N*_*e*_ obtained by splitting the X chromosome into six pieces to enable bootstrapping with the *N*_*e*_ estimated on the undivided X chromosome, and the results are consistent (Figure S3).

We compared *N*_*e*_ estimated from the simulated autosome and X chromosome data to the actual *N*_*e*_ for the UK-like demographic model. The actual autosome or X chromosome *N*_*e*_ can be obtained from the sex-specific effective population sizes using Equations 2 and 3. The estimated *N*_*e*_ generally matches the true *N*_*e*_ closely (Figure 1 and Figure S4). However, some discrepancies exist between the estimated and actual *N*_*e*_ because IBDNe cannot localize sharp changes in the population size to one exact generation and it tends to over-smooth corners of the trajectory of effective population-size over time (Browning and Browning 2015).

**Figure 1.**
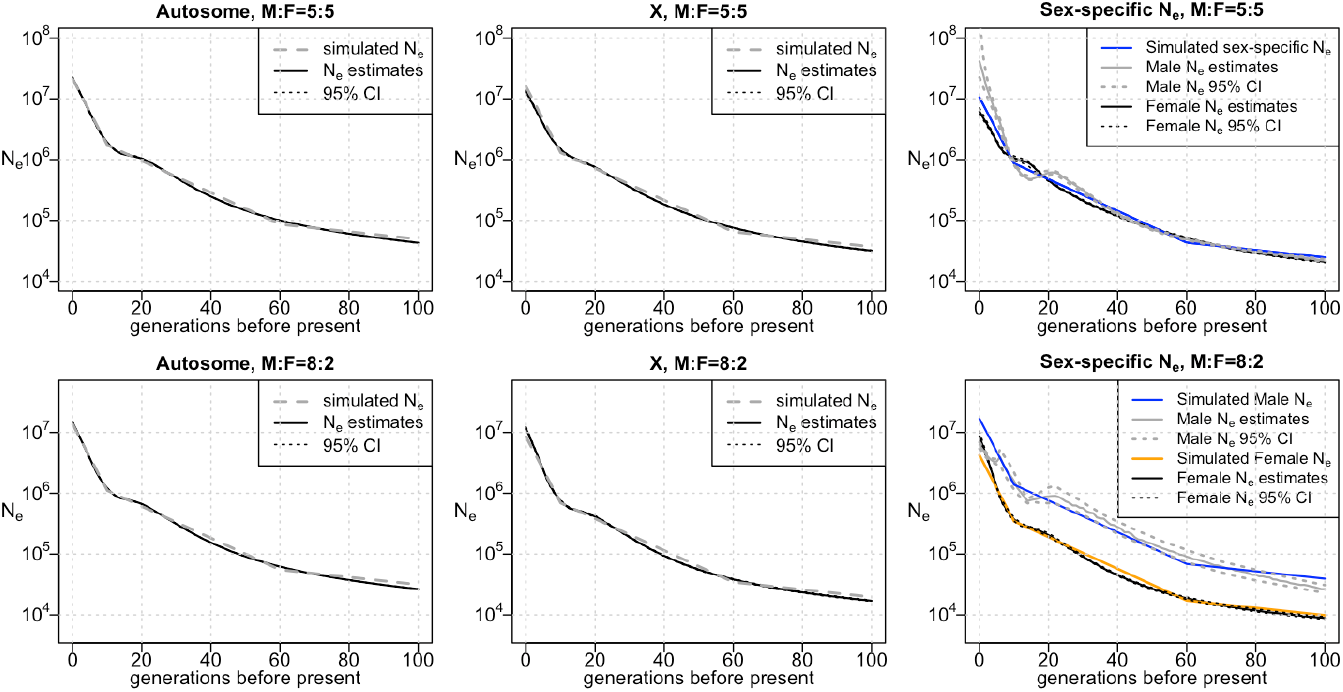
Estimates of autosome, X chromosome, and sex-specific *N*_*e*_ in UK-like simulation study. Autosomal *N*_*e*_ is shown in the left column, X chromosome *N*_*e*_ in the middle column, and sex-specific *N*_*e*_ in the right column. The top row is with equal sex ratio, while the second row is with 20% females. Results for other choices of sex-ratio can be found in Figure S4. Y-axes show *N*_*e*_ plotted on a log scale. In cases where the estimated sex-specific *N*_*e*_ is negative (see Methods), it is not shown.

We next used the estimated X chromosome *N*_*e*_ and autosome *N*_*e*_ to estimate the sex-specific effective population sizes. We find that the formulas for the sex-specific *N*_*e*_ as functions of the autosome and X chromosome *N*_*e*_(Equation 5, 6, 7) are sensitive to errors in *N*_*e*_ estimation. The estimated sex-specific *N*_*e*_ are similar to the actual values most of the time, but at around 15 generations ago, where there was a large change in the population growth rate, we observe greater inaccuracy in the estimated *N*_*e*_ and the estimated sex-specific *N*_*e*_ differs from the actual sex-specific *N*_*e*_ (Figures 1 and Figure S4).

### UK Biobank data

The estimated autosome *N*_*e*_ for the UK Biobank’s White British population (Figure 2) went through a period of moderate growth between 50 and 100 generations ago. Between 20 and 50 generations ago the effective population size was fairly constant. The effective population size had a high rate of growth in the most recent 20 generations and reached a current population size of 169 million (95% confidence interval = 139-221 million). IBDNe estimates the most recent generations by extrapolating the growth rate of earlier generations and doesn’t account for a possible recent decrease in population growth rate, hence the *N*_*e*_ for generation 0 may be overestimated (Browning and Browning 2015). The autosome *N*_*e*_ estimated from the genotype data of the UK Biobank Indian participants (Figure 3) shows slow growth from 100 generations ago until around 10 generations ago and a higher rate of growth in the past 10 generations. The estimated current effective population size is 89 million (95% confidence interval = 47-221 million).

**Figure 2.**
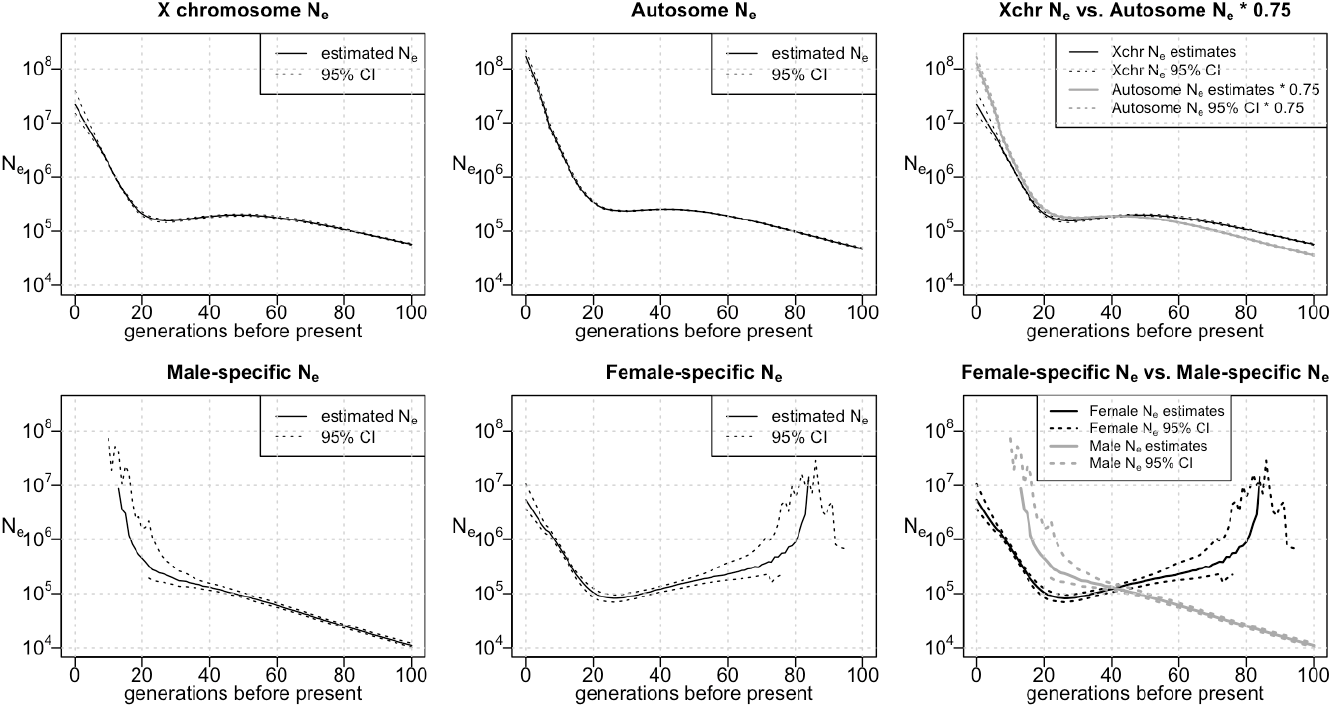
Effective population size of the UK Biobank White British group. From left to right, the top row shows the estimated X chromosome *N*_*e*_, the estimated autosome *N*_*e*_, and a comparison of the estimated X chromosome *N*_*e*_ with 75% of the estimated autosome *N*_*e*_. The bottom row displays the male-specific *N*_*e*_, the female-specific *N*_*e*_, and a comparison between them. For the *N*_*e*_ plots, the Y-axes show *N*_*e*_ on a log scale. In cases where the estimated sex-specific *N*_*e*_ or its confidence band is negative (see Methods), the negative values are not shown.

**Figure 3.**
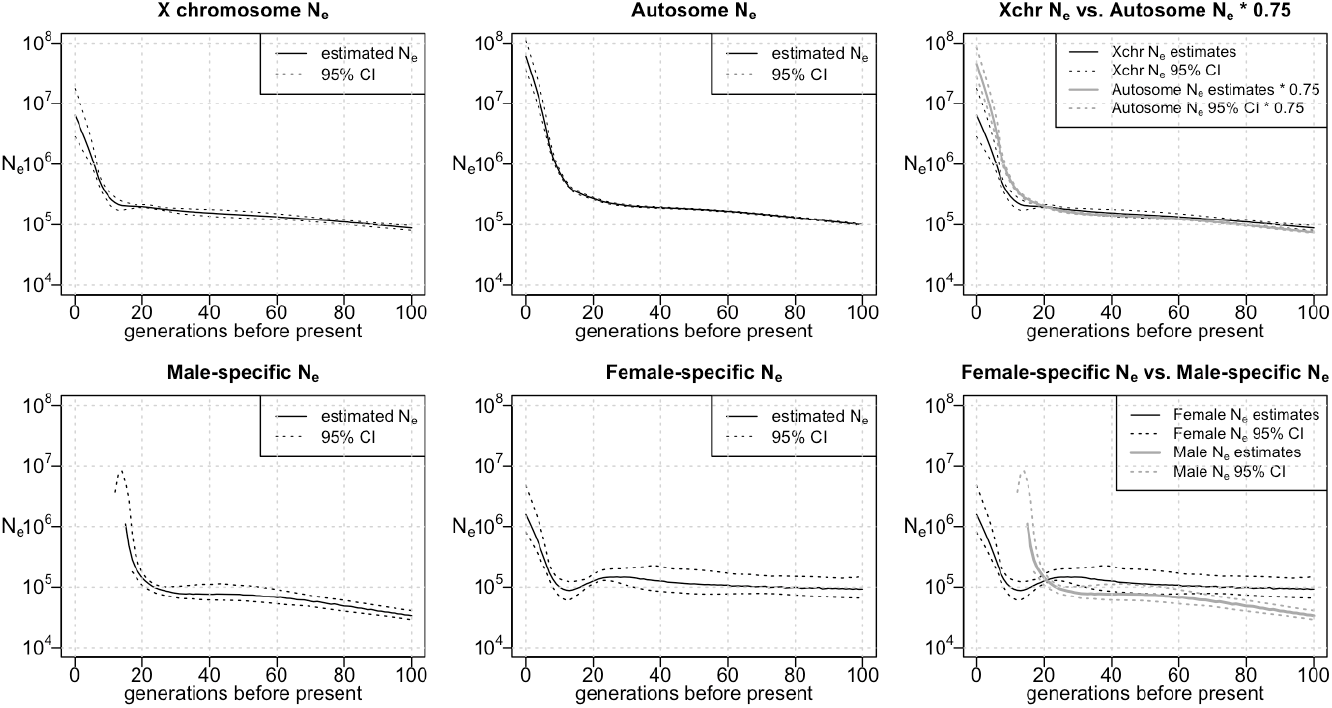
Effective population size of the UK Biobank Indian group. From left to right, the top row shows the estimated X chromosome *N*_*e*_, the estimated autosome *N*_*e*_, and a comparison of the estimated X chromosome *N*_*e*_ with 75% of the estimated autosome *N*_*e*_. The bottom row displays the male-specific *N*_*e*_, the female-specific *N*_*e*_, and a comparison between them. For the *N*_*e*_ plots, the Y-axes show *N*_*e*_ on a log scale. In cases where the estimated sex-specific *N*_*e*_ or its confidence band is negative (see Methods), the negative values are not shown.

The inferred X chromosome *N*_*e*_ has a similar shape to the inferred autosome *N*_*e*_ for both groups. Notably, the estimated X chromosome *N*_*e*_ is fairly close to 75% of the inferred autosomal effective population size, which is the expected effective population size that would be obtained from the X chromosome data if the female and male effective population sizes have been equal in these populations (Figures 2 and 3). However, the estimated 75% autosome effective size is lower than the X chromosome effective size between 42 and 100 generations ago for the UK Biobank White British, while the estimated 75% autosome effective size is higher than the X chromosome effective size in the past 41 generations although the confidence intervals for the two estimates overlap in parts of this range. In the UK Biobank Indian group we see a similar pattern, with higher estimated X chromosome size between 22 and 100 generations ago and higher estimated 75% autosome effective size in the past 21 generations, although in the period between 22 and 100 generations ago the estimates are very close and the confidence intervals largely overlap.

To investigate the historical sex-specific effective population sizes of the UK Biobank White British population and the UK Biobank Indian population, we used the X chromosome and autosome *N*_*e*_ to estimate the female and male *N*_*e*_ of these two populations. For the UK Biobank White British, there is an apparent dip in female effective size around 27 generations ago (Figure 2, bottom center), but this may be an artifact similar to the dip in estimated effective size in the simulated UK-like population around 15 generations ago (Figure 1, top right); both of these dips occur around the timing of an increase in population growth rate. For the Indian group, a similar dip appears in estimated female effective size around 13 generations ago. For both groups, ignoring generations for which estimates are negative, we find that in the more recent past (up to 41 generations ago for the UK Biobank White British group and up to 21 generations ago for the UK Biobank Indian group) the estimated male effective population size is higher than the female, while this is reversed in the more distant past.

The sex-specific *N*_*e*_ estimates for both the White British group and the Indian group contain negative values. The estimated female effective population size of the White British group is negative earlier than 86 generations before the present due to overly high X chromosome *N*_*e*_ estimates compared to autosome *N*_*e*_ estimates. Likewise, the estimated male effective population size of this group is negative from 13 generations ago to the present as the X chromosome *N*_*e*_ estimates is low compared to autosome *N*_*e*_ estimates. In addition, the estimated male effective population size of the Indian group is negative from 13 generations ago to the present as a result of excessively low X chromosome *N*_*e*_ estimates compared to autosome *N*_*e*_ estimates. The negative *N*_*e*_ estimates over these periods are not scientifically meaningful and are generally accompanied by large confidence intervals. For both the White British group and the Indian group, the distribution of X-chromosome IBD lengths are mostly consistent in male-male, female-male, or female-female haplotypes. The rate of IBD segments that are longer than 8 cM (sex-averaged unit) is slightly higher for male-male IBD haplotypes in the Indian group (Figure S2). However, the slight difference among rates of long IBD segments in the three categories is likely a consequence of sampling variability as discussed in the legend of Figure S2.

## Discussion

Previous studies have shown that the X chromosome can provide information about demographic processes that cannot be revealed by the analysis of autosomes alone (Ramachandran *et al*. 2004; Schaffner 2004; Bustamante and Ramachandran 2009; Bryc *et al*. 2010; Goldberg and Rosenberg 2015; Shringarpure *et al*. 2016). In this work, we focused on utilizing IBD segments on the X chromosome to infer the X chromosome effective population size. We developed a framework to model X chromosome *N*_*e*_ and derived the relationship between X chromosome and autosome *N*_*e*_ by considering the different coalescence rates between X chromosomes and between autosomes. We also showed how to apply this information to calculate the female and male effective population sizes in a population as functions of the X chromosome and autosome *N*_*e*_.

We applied our method to estimate the X chromosome effective population size for the UK Biobank White British individuals and the UK Biobank Indian individuals. In both populations, we observe a time point within the past decades of generations when male estimated *N*_*e*_ intersected with and continued to exceed female estimated *N*_*e*_. We speculate that demographic trends such as polygamy that favored a lower male effective population size may have been more prevalent in the more distant past, and that migration rates, which have increased in recent times, may have increased more in males than in females which would increase the male effective population size relative to the female effective population size. However, it is also possible that artifacts such as better detection of long IBD segments on the X chromosome due to more accurate phasing resulting from the smaller effective population size and haploid males could be responsible for these trends.

We validated the performance of our method in a simulated population with similar growth rates and IBD rates as the UK population. We found that the estimated sex-specific effective population sizes capture the overall trends of the true female and male effective population sizes. In the simulations, however, we observe some significant discrepancies between the estimated sex-specific *N*_*e*_ and the true sex-specific *N*_*e*_ across timescales of tens of generations, which tends to occur around points in time at which the population’s growth rate changed. This indicates that the estimated sex-specific *N*_*e*_ calculated from the estimated autosomal and X chromosome *N*_*e*_ should not be used to test hypotheses about differences in effective population size between the sexes, particularly when those hypotheses involve differences that occur for limited time periods rather than across the full estimation timescale.

Although X chromosome IBD information has been used by previous studies for the estimation of genealogical relations between individuals, especially for kinship estimation in forensic settings (Pinto *et al*. 2011; Buffalo *et al*. 2016; Henden *et al*. 2016), there has been a lack of studies that use X chromosome IBD segments to estimate recent effective population size. Our work thus fills a gap that existed in the application of X chromosome IBD information in population genetic studies.

Previous methods for estimating sex-specific population history or sex bias in human populations have relied on comparisons of genetic diversity between autosomes and the X chromosome using allele frequency differentiation, patterns of neutral polymorphism, and the site frequency spectrum (Ramachandran *et al*. 2004; Hammer *et al*. 2008; Keinan *et al*. 2009; Emery *et al*. 2010; Clemente *et al*. 2018; Musharoff *et al*. 2019). Most of these methods considered only a single estimate of the effective sex ratio over the entire history of a population, although this ratio can vary over time (Hammer *et al*. 2008; Keinan *et al*. 2009; Emery *et al*. 2010; Clemente *et al*. 2018). Some of these methods also focused on a constant overall effective population size across time, although changes in effective population size can distort these analyses (Ramachandran *et al*. 2004; Keinan *et al*. 2009). Recently, Musharoff et al. developed a likelihood ratio test for population sex bias that considered populations of non-constant size and changing sex ratios using site frequency spectrum data (Musharoff *et al*. 2019). However, this method requires demographic parameters to be constant within time epochs. In comparison, our approach for estimating the X chromosome effective population size and the sex-specific effective population sizes requires minimal assumptions and allows the effective population sizes to vary independently over time. The ability of our IBD-based analyses to infer effective population sizes in the past hundred generations distinguishes our approach from other methods.

There are several limitations to our approach. First, IBD-based estimation of effective population size requires a large sample of individuals from the population. The performance of IBDNe is affected by the number of detected IBD segments. For example, we observe wider confidence bands for the estimated *N*_*e*_ in the most recent generations since there tend to be fewer very long IBD segments in the sample. Similarly, we observe wider confidence bands for the X chromosome *N*_*e*_ compared to the autosomal *N*_*e*_ due to the smaller amount of data in the X chromosome.

Second, our method for estimating *N*_*e*_ is less accurate around times when the population experienced an abrupt change in population size or growth rate. It tends to over-smooth the autosomal and X chromosome trajectories, which can then induce spurious oscillations in the estimated sex-specific effective population size over time. Moreover, these artifacts persist in the bootstrapped data, so that the bootstrap confidence intervals are over-confident in such regions and do not have the desired level of coverage of the underlying effective population sizes.Given these limitations, we note that although our method provides reliable estimates of the X chromosome effective population size in simulated data, its application to estimate sex-specific population history is not suitable for rigorously testing hypotheses on sex-specific *N*_*e*_ in a population, because small inaccuracies in estimation of autosome and X chromosome effective population sizes are magnified when transforming these to estimates of sex-specific effective population size. We thus recommend only considering the estimated sex-specific effective population size as a tool to explore the overall pattern of sex-specific past demographic events.

## Data and code availability

UK Biobank data were obtained via the UK Biobank’s Research Analysis Platform. The IBDNe software is available from https://faculty.washington.edu/browning/ibdne.html. The hap-ibd software is available from https://github.com/browning-lab/hap-ibd.

## Acknowledgements

Research reported in this publication was supported by the National Human Genome Research Institute of the National Institutes of Health under award number HG005701. The content is solely the responsibility of the authors and does not necessarily represent the official views of the National Institutes of Health.

This research has been conducted using the UK Biobank Resource under Application Number 19934.

## Supplementary Figures

**Figure S1.**
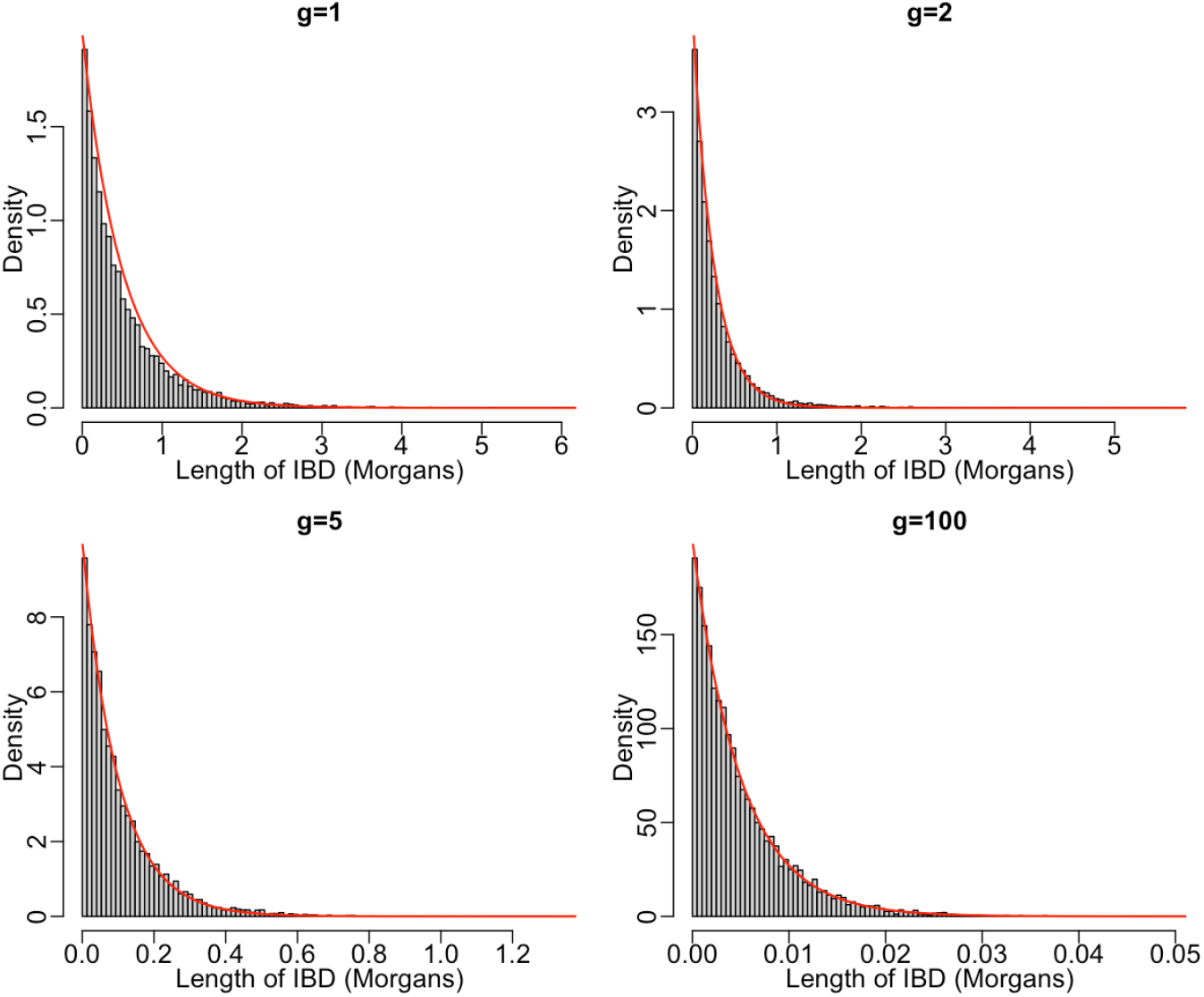
Simulated distribution of the length of an X chromosome IBD segment measured in sex-averaged genetic distance given the number of generations to the shared ancestor. The distribution of 10,000 simulated exponential(3*F*/2) observations, where *F*∼binomial(2*g*, 2/3), is compared to the exponential(2*g*) density, shown as the red curve in each plot, for *g* = 1, 2, 5 and 100 generations. The simulated distribution is for the distribution of the length of the segment when assuming Haldane’s model, while the exponential(2*g*) is the distribution used in our methodology. When *F* = 0, the simulated length of IBD segment is set to 100 Morgans. Although not shown in the figure, the observations corresponding to *F* = 0 were accounted for when calculating the simulated density.

**Figure S2.**
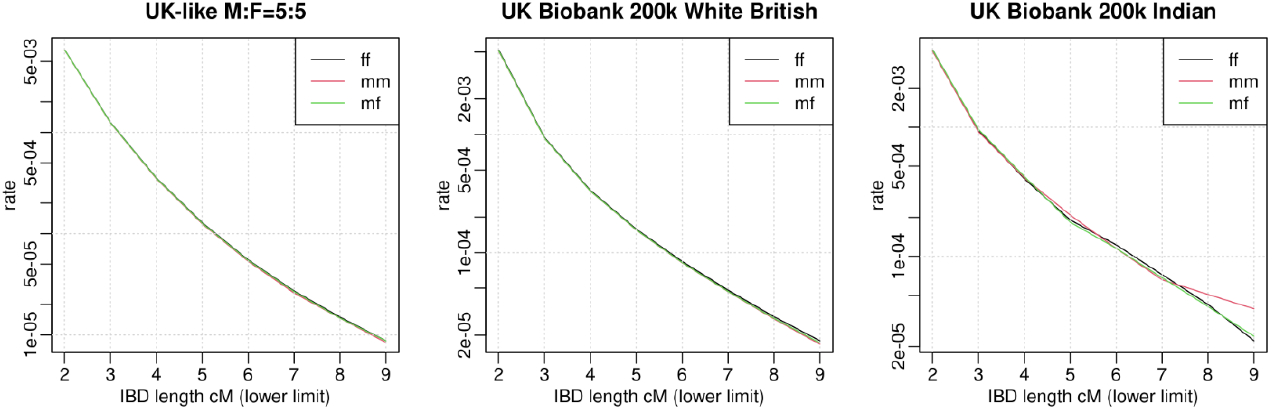
The distribution of X chromosome IBD segments between sexes. The black line, red line and green line show the rate of IBD segments on a log scale over a series of consecutive length bins in female-female (ff) haplotype pairs, male-male (mm) haplotype pairs, and male-female (mf) haplotype pairs, respectively. The rate of IBD segments in each length bin for each sex combination is calculated as the number of IBD segments from pairs of individuals with the corresponding sex in that bin divided by the total number of haplotype pairs of the corresponding sex combination. IBD lengths are measured in sex-averaged units. The left column displays results from the UK-like simulation with equal sex ratio. The middle column displays results from the White British group in the UK Biobank sequence data. The right column displays results from the Indian group in the UK Biobank sequence data. There is a slightly higher rate of male-male IBD segments that are longer than 8 cM (sex-averaged unit) in the Indian group, but this difference may not be significant given the small number of IBD segments in the three categories (204 in female-female haplotypes, 71 in male-male haplotypes, and 205 in male-female haplotypes). Since the number of IBD segments with length in this range are approximately Poisson distributed, the magnitude of the difference observed between the count of male-male haplotypes and female-female or female-male haplotypes is about 2x pooled standard deviations.

**Figure S3.**
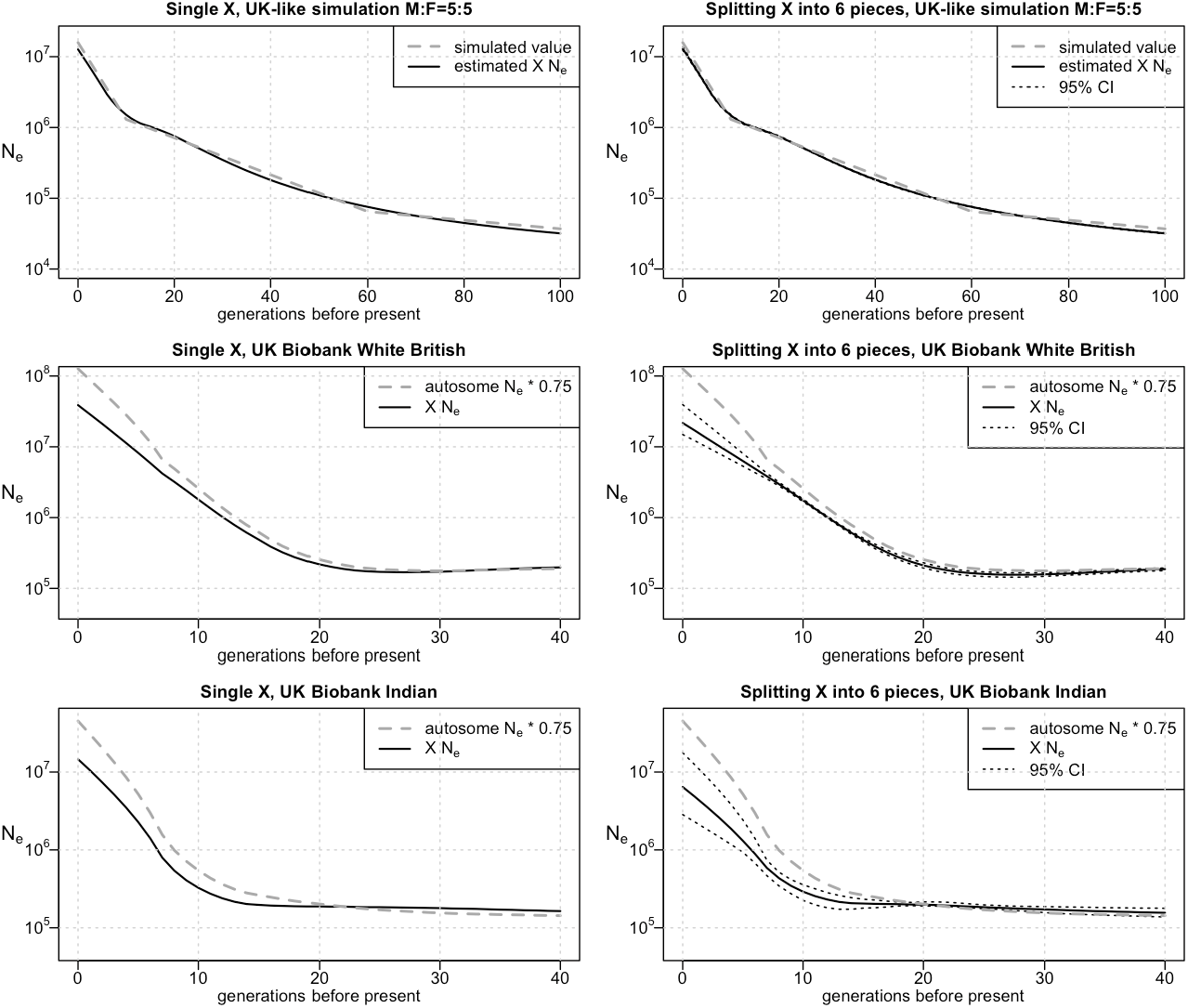
Effective population size estimated from an undivided X chromosome versus the effective population size estimated by splitting the X chromosome into six regions. *N*_*e*_ estimated on a single undivided X chromosome is shown in the left column. *N*_*e*_ estimated by treating six separate regions of the X chromosome as six chromosomes to enable bootstrapping is shown in the right column. In each plot, the Y-axes are on a log scale. From top to bottom, the rows display results from a UK-like simulation with equal sex ratio, the White British group in UK Biobank, and the Indian group in UK Biobank.

**Figure S4.**
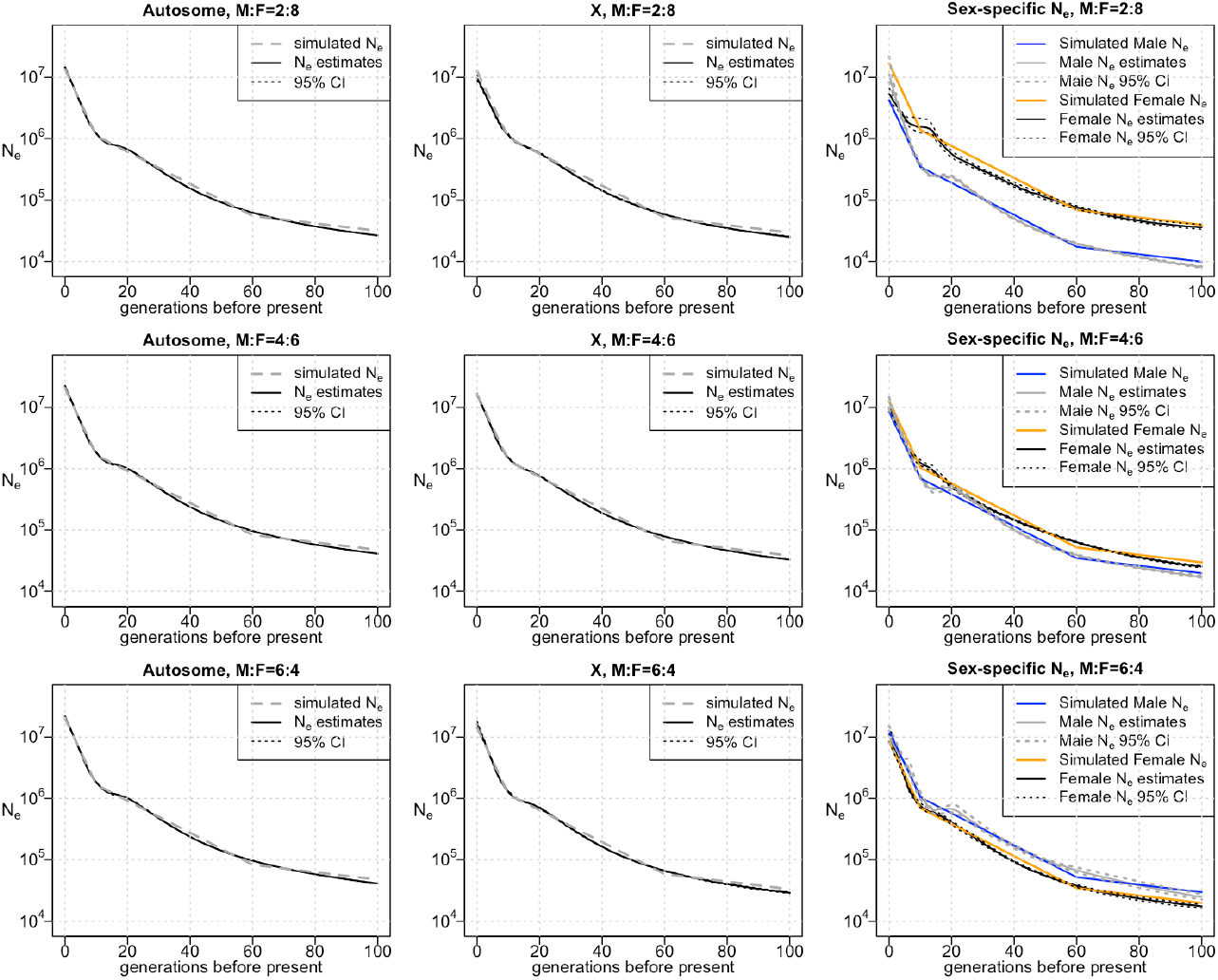
Estimates of autosome, X chromosome, and sex-specific *N*_*e*_ in UK-like simulations. Autosomal *N*_*e*_ is shown in the left column, X chromosome *N*_*e*_ in the middle column, and sex-specific *N*_*e*_ in the right column. From top to bottom, the rows display results from a UK-like simulation with 80%, 60%, and 40% females. The Y-axes are on a log scale.

